# Reconstruction of the Prox gene family evolution in vertebrates reveals multiple lineage-specific gene losses

**DOI:** 10.64898/2026.09.25.754218

**Authors:** Virginia Panara, Jake Leyhr, Katarzyna Koltowska, Tatjana Haitina

## Abstract

Prospero-related homeobox (Prox) genes encode a family of transcription factors that play essential roles in the development of several organs and systems, including the central nervous system, lymphatic endothelium, musculature, and liver. Despite their developmental importance, the evolutionary history of the vertebrate Prox gene family remains poorly understood.

In this study we combined phylogenetic and synteny analysis to characterise the evolution of the Prox family in vertebrates. Our results reveal that two to three Prox subfamilies were already present in the last common ancestor of jawed vertebrates. We clarify the identity and evolutionary relationships of well-studied members of this family and identify multiple independent losses of Prox2 and Prox3 genes in specific vertebrate lineages. Furthermore, we uncover evidence for the existence of a fourth Prox gene in the ancestral vertebrate genome, which was subsequently lost.

Overall, this study provides the first comprehensive analysis of the evolutionary history of the vertebrate Prox gene family and establishes a foundations for future studies on the functional roles of these genes.

## Introduction

Prospero-related homeobox (Prox) genes are a family of transcription factors (TFs) playing crucial roles in the development of several key organs and systems. *Prox* genes are part of the larger PROS class, which is characterised by the presence of an atypical homeodomain with DNA binding activity (Elsir et al. 2012), and have been connected to functions such as positive and negative regulation of gene expression (Kivelä et al. 2016; Zhang et al. 2017), exit from the cell cycle and differentiation (Kaltezioti et al. 2010), and cell migration (Mishima et al. 2007). The vertebrate members of this family are divided into three subfamilies: Prox1, Prox2 and Prox3. Prox1 is by far the most studied member, with well-described roles in the development of the retina (Dyer et al. 2003), lens (Wigle et al. 1999), liver (Sosa-Pineda et al. 2000; Oliver et al. 1993), pancreas (Wang et al. 2005; Oliver et al. 1993; Burke and Oliver 2002), muscles (Oliver et al. 1993; Kivelä et al. 2016), central nervous system (Torii et al. 1999; Lavado and Oliver 2007), peripheral nervous system (Becker et al. 2010; Holzmann et al. 2015), lymphatic vessels (Wigle and Oliver 1999; Nicenboim et al. 2015; Koltowska et al. 2015) and heart (Risebro et al. 2009; Kivelä et al. 2016). The other two members of the family haven’t been characterised to the same extent, although recently *Prox2* has been studied for its role in the development of the sensory vagal neurons (Lowenstein et al. 2023).

Despite their important roles in development, little attention has been paid to the phylogenetic reconstruction and classification of the members of the *Prox* family. For example, zebrafish *prox1b* has been extensively characterised for its expression and function in the lymphatic vasculature (Giacco et al. 2010; Tao et al. 2011; van Impel et al. 2014; Koltowska et al. 2015; Grimm et al. 2023). However, the gene identity was never properly established by phylogenetic analysis in these publications (Giacco et al. 2010), and later studies place it as part of a separate *Prox* clade (Lanoizelet et al. 2024). Therefore, an in-depth classification of the members of the *Prox* family is needed in order to reconstruct the evolutionary history of the *Prox* family and correctly interpret the available data.

Phylogenetic analysis is the main tool used to assign gene identities and reconstruct the evolutionary history of gene families. However, synteny (the analysis of the conserved gene location along a chromosome) can support and complement the results of phylogeny, helping in resolving complex cases such as those arising through the reciprocal loss of part of the same gene family in related clades. For example, a combined phylogenetic-syntenic approach has been used to elucidate, among others, the evolutionary history of the NK4 (Ford et al. 2025), the carbohydrate 6-O sulfotransferase (Ocampo Daza and Haitina 2020), the corticotropin- releasing hormone (Cardoso et al. 2020), and the protein kinase C (Garcia-Concejo and Larhammar 2021) gene families.

In this study, we applied a combined phylogenetic and a synteny-based approach to reconstruct the evolutionary history of the *Prox* family in vertebrates. Our analyses revealed that the *Prox1*, *Prox2,* and *Prox3* subfamilies were already present in the last common ancestor of vertebrates and Prox2 and Prox3 genes were lost in specific lineages. Moreover, we reconstructed the presence of a fourth *Prox* clade in vertebrates that has been lost in all extant groups. Finally, we propose a model for Prox gene functional evolution in vertebrates, whereby an ancestral function in the central nervous system was retained and expanded to other organs by the *Prox1* and *Prox3* subfamilies, possibly leading to a functional redundancy, potentially contributing to the repeated loss of *Prox3* in several lineages. Overall, this study provides the first in-depth characterization of the *Prox* family evolutionary history in vertebrates and lays the foundation for a better understanding the roles of these genes in development across vertebrates.

## Results

### Three distinct Prox subfamilies are present in gnathostomes

In order to establish the identity of *Prox* genes and their relation to each other, we performed a phylogenetic analysis. The resulting tree (**Figure 1A**) was rooted using the invertebrate species. The tree recovered three distinct Prox gene subfamilies in jawed vertebrates (or gnathostomes): Prox1, Prox2 and Prox3 (**Figure 1A-B**). The analysis was repeated using a different alignment method (MUSCLE), obtaining the same relation between the three Prox clades (**Figure S1A, S1B**). The Prox2 clade is sister to the other two and includes sequences from all major gnathostome groups (**Figure 1B**). Several of the *Prox2* sequences were annotated as *Prox1* or *Prok1-like*, including the *Prox2* genes in *Protopterus annectens, Lates calcarifer, Acipenser ruthenus,* and *Amia calva*. In a few of the considered teleost species, such as *Takifugu flavidus*, *Lates calcarifer,* and *Oreochromis niloticus*, two copies of *Prox2* could be identified. The presence of two *Prox2* genes was further confirmed by BLAST on the GenBank database in several additional teleost species, including *Pundamilia nyererei, Channa argus, Anarhichas minor,* and *Cyclopterus lumpus*, suggesting the two teleost copies came from the teleost 3R whole genome duplication (WGD), and that one copy was later lost in species such as zebrafish.

**Figure 1:**
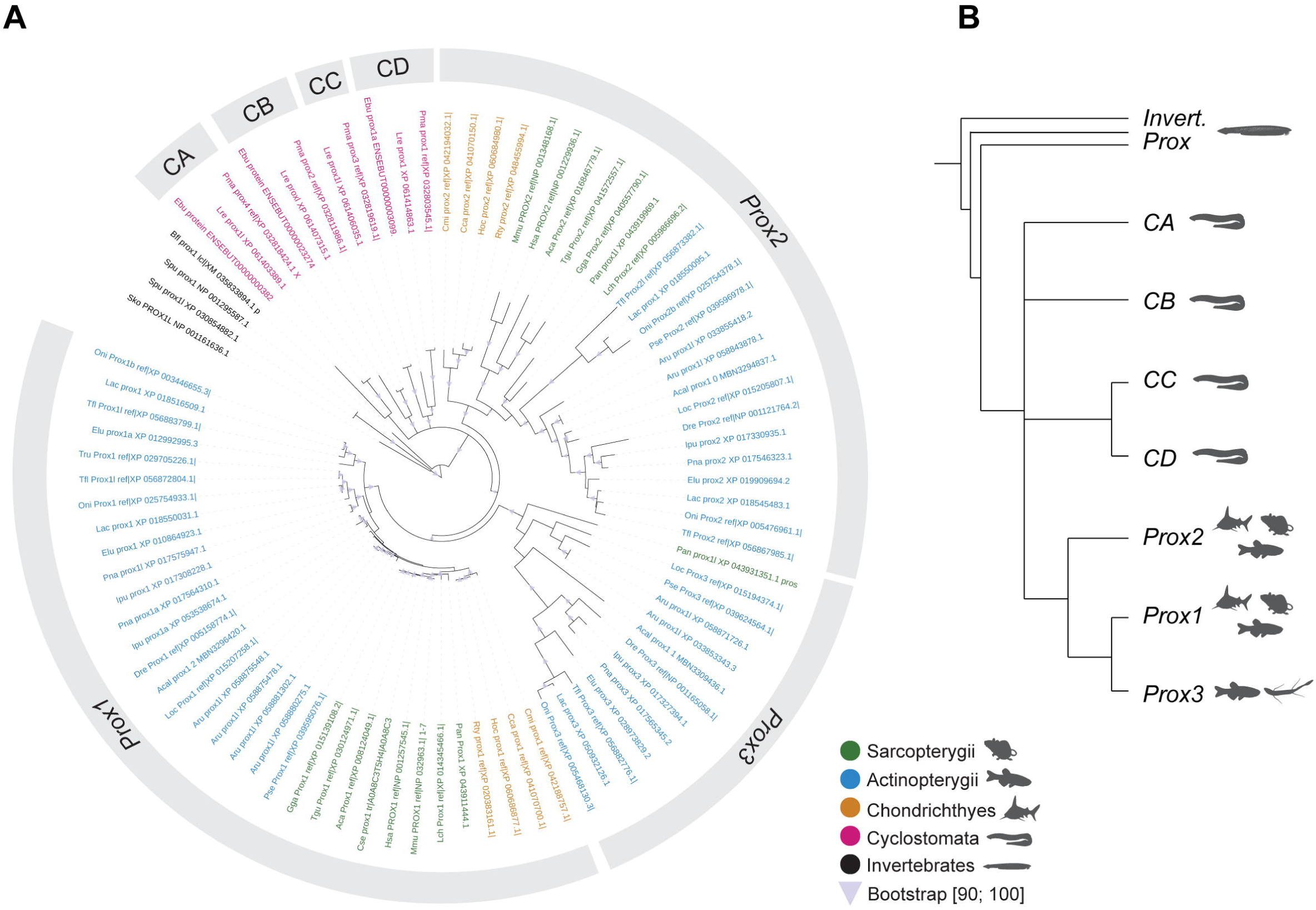
Phylogenetic analysis of the vertebrate *Prox* family. A) Consensus tree of the *Prox* family protein sequences based on the Clustal Omega alignment. Nodes with bootstrap value < 50 have been collapsed. Triangles: nodes with [90:100] bootstrap values. Green: sarcopterygian sequences. Blue: actinopterygian sequences. Yellow: chondrichthyan sequences. Magenta: cyclostome sequences. Black: invertebrate/other sequences. B) Schematic cladogram of the *Prox* family established in A.

*Prox1* is present throughout gnathostomes, and its members have shorter branch lengths compared to other clades (**Figure 1A**), consistent with its conserved developmental role (Elsir et al. 2012) and the previously observed high degree of conservation of its enhancer landscape (Panara et al. 2024). Following the teleost 3R, several species retain two copies of *Prox1*, including *Takifugu flavius, Oreochromis niloticus,* and *Lates calcarifer* (**Figure S2B**). However, in zebrafish, only one copy of *Prox1* (*prox1a*) can be found based on phylogeny, and the second copy has been lost.

Sister to *Prox1* is the *Prox3* clade, mainly composed of actinopterygian sequences. The presence of a single sarcopterygian sequence in lungfish (*Protopterus annectens*) in the *Prox3* clade, further confirmed by synteny analysis (**Figure S2D**), dates the split between *Prox1* and *Prox3* to the ancestor of Osteichthyes. This presence also suggests the gene has been lost in the tetrapod lineage, and possibly also independently lost in the coelacanth lineage, unless we were unable to identify it in the coelacanth genome due to some remaining genome assembly gap.

The *Prox3* clade also includes a zebrafish gene, previously described as zebrafish *prox1b*, a co-ortholog of *prox1a* arising from the teleost 3R (Giacco et al. 2010). Instead, this gene belongs to the *Prox3* clade and appears to have arisen from a much earlier duplication event. For this reason, the gene name has been changed to *Dre_prox3* in **Figure 1A** to reflect its correct identity. Only one copy of *prox3* was identified in the teleost species used for the phylogenetic analysis. The presence of a single *prox3* gene in teleosts was confirmed by reciprocal BLAST on the GenBank database. Additional copies of *prox3* could only be detected in species known to have undergone additional lineage specific WGD events, such as *Cyprinus carpio* (Xu et al. 2014), *Carassius auratus* (Chen et al. 2019), *Myxocyprinus asiaticus* (X. Liu et al. 2022),

*Sinocyclocheilus anshuiensis* (Li et al. 2021), *Coregonus clupeaformis* (Allendorf and Thorgaard 1984; Sutherland et al. 2016), and *Oncorhynchus clarkii lewisi* (Allendorf and Thorgaard 1984). We could not identify any *prox3* genes in the chondrichthyan species considered in the phylogeny (**Figure 1A**). An additional search by reciprocal BLAST on the GenBank databases for holocephalans, sharks, rays, and skates also failed in identifying any *Prox3* gene in chondrichthyans. The presence of both *Prox1* and *Prox2* in this clade suggests the *Prox3* was independently lost in this group.

Therefore, we can confirm that three *Prox* gene subfamilies are present in jawed vertebrates. We also determined that *Prox3* has been independently lost in several lineages and includes the gene previously named *prox1b* in zebrafish.

### Local synteny reveals a conserved gene landscape surrounding Prox genes

In the phylogenetic analysis, cyclostome *Prox* sequences from two lamprey species (*Lethenteron reissneri* and *Petromyzon marinus*) and one hagfish species (*Eptatretus burgeri*) were considered. Four *Prox* genes were identified for both lamprey species, and three for the hagfish. Four cyclostome gene clades (named CA-CD in **Figure 1A-B**) were found in a polytomy (CA, CB, CC/CD) at the base of the vertebrate *Prox* clade. This phylogenetic positioning is likely due to low-support for any specific tree topology, a well-documented problem when working with cyclostome sequences (Kuraku et al. 2008). We attempted to determine the position of these clades by re-running the phylogenetic analysis, including only one of the at the time at a the time, but this did not result in a clearer placement (**Figure S2A**).

In order to correctly identify the cyclostome Prox genes local synteny was employed, as this approach can be used in combination with phylogenetic analysis to assign the identity of genes. To investigate the local synteny surrounding the *Prox* genes, we developed the EzGeneSynteny Python package to automatically download the names and relative orientations of the loci neighbouring our sequences of interest. Briefly, EzGeneSynteny queries NCBI GenBank via Entrez to retrieve the genomic region surrounding each input gene and extracts upstream and downstream annotated coding sequences, generating simple gene order maps to assist in confirming gene orthology assignments across species. BLAST against GenBank and Ensembl was used to verify the EzGeneSynteny output. Gene families with members in at least two of the identified clades (Prox1-3, CA-D, and invertebrate Prox) were further considered for this analysis and plotted in **Figure S2B-J**.

Within gnathostomes, a high level of conserved synteny was found within each *Prox* subfamily. E.g., the position of *Dlst* and *Snx15* upstream of *Prox2* and *Prox3,* respectively, is conserved in all the gnathostome species considered (**Figure S2C-D**). Overall, these patterns of conserved synteny support the three gnathostome clades identified by the phylogenetic analysis.

The analysis also identified elements of the ancestral syntenic organisation. For example, the placement of a *Naalad/Naaladl* locus downstream of *Prox* can be observed both in the *Prox3* clade and in the lancelet (*Branchostoma floridae*) *Prox* gene (**Figure S2D, S2J**), suggesting *Naalad/Naaladl* was present in the ancestral locus before the divergence of chordate and vertebrate lineages. Similarly, the placement of *Dlst* upstream of *Prox* is a feature conserved in chordates and even Ambulacraria (**Figure S2C-S2J**), implying this gene has retained its position with respect to *Prox* since the most recent common ancestor of deuterostomes.

### Rps6kc1, Rps6kl1, and Snx15 paralogs can be used to further reconstruct the Prox clade phylogeny

The position of *Rps6kc1* and *Rps6kl1* upstream of *Prox1* and *Prox2*, respectively, is conserved in jawed vertebrates (**Figure 2A**, **S2B-C**). By reciprocal BLAST, we could determine that the *Snx15* locus upstream of *Prox3* (**Figure 2A**, **S2D**) is also closely related to *Rps6kc1* and *Rps6kl1,* despite the different nomenclature. Using the Uniprot feature viewer (https://www.uniprot.org) and Uniparc, we determined that *Snx15* shows high conservation with the N-terminal portion of *Rps6kc1*, including a PX protein domain (**Figure 2B**), while *Rps6kl1* shows conservation with the C-terminal portion of *Rps6kc1*, including the presence of a protein kinase domain (**Figure 2B**). To confirm these observations, the mouse *Snx15* protein sequence (NP_081188.1) was used as the query for a BLASTP search, which returned *Rps6kc1* as a higher hit than any other members of the *Snx* family. Moreover, the Human Gene Database and Ensembl Paralogue page of human *SNX15* list *RPS6KC1* and *RPS6KL1* as paralogs. These loci, positioned upstream of each of the gnathostome *Prox* clade, appear to be closely related to each other, making them particularly interesting for the reconstruction of conserved synteny. Moreover, these genes are found upstream of the cyclostome *Prox* genes, and can therefore be leveraged for determining their identity by syntenic analysis. Therefore, a phylogenetic analysis of the *Rps6k* family was run. A BLASTP search was used to identify the relevant sequences (**Table S2**). In lancelet, the two retrieved hits, named *Snx15* and *Rps6kl1*, are located in tandem on chromosome 1 and are not in close proximity to a *Prox* locus. Phylogenetic analysis revealed that *Bfl Snx15* and *Rps6kl1* group with vertebrate *Snx15* and *Rps6kl1* sequences, respectively (**Figure S3A-B**). However, a close inspection revealed that the last three exons of *Snx15* are also annotated as part of *Rps6kl1* (**Figure 2C**). We therefore hypothesise this is an annotation mistake and combined the two sequences into a single coding sequence. We verified the new sequence matches the domain structure of *Rpskl1* by alignment, which identified the three conserved protein domains (PX, MIT, and PK) in the expected order and position. Apart from preventing skewing in the analysis, this also allowed us to assign this one lancelet gene as the outgroup to the rest. For the phylogenetic analysis, sequences were aligned in Seaview using the Clustal Omega method, as MUSCLE gave very suboptimal alignments. The resulting tree retrieved the *Rps6kl1* clade as sister to the rest (**Figure 2D**). Since *Rps6kl1* is found in synteny with *Prox2*, the *Rps6k* family tree topology matches the *Prox* one, which places *Prox2* as sister to the other *Prox* genes (**Figure 2E-F**). The results from this analysis were incorporated in the following local synteny analysis.

**Figure 2:**
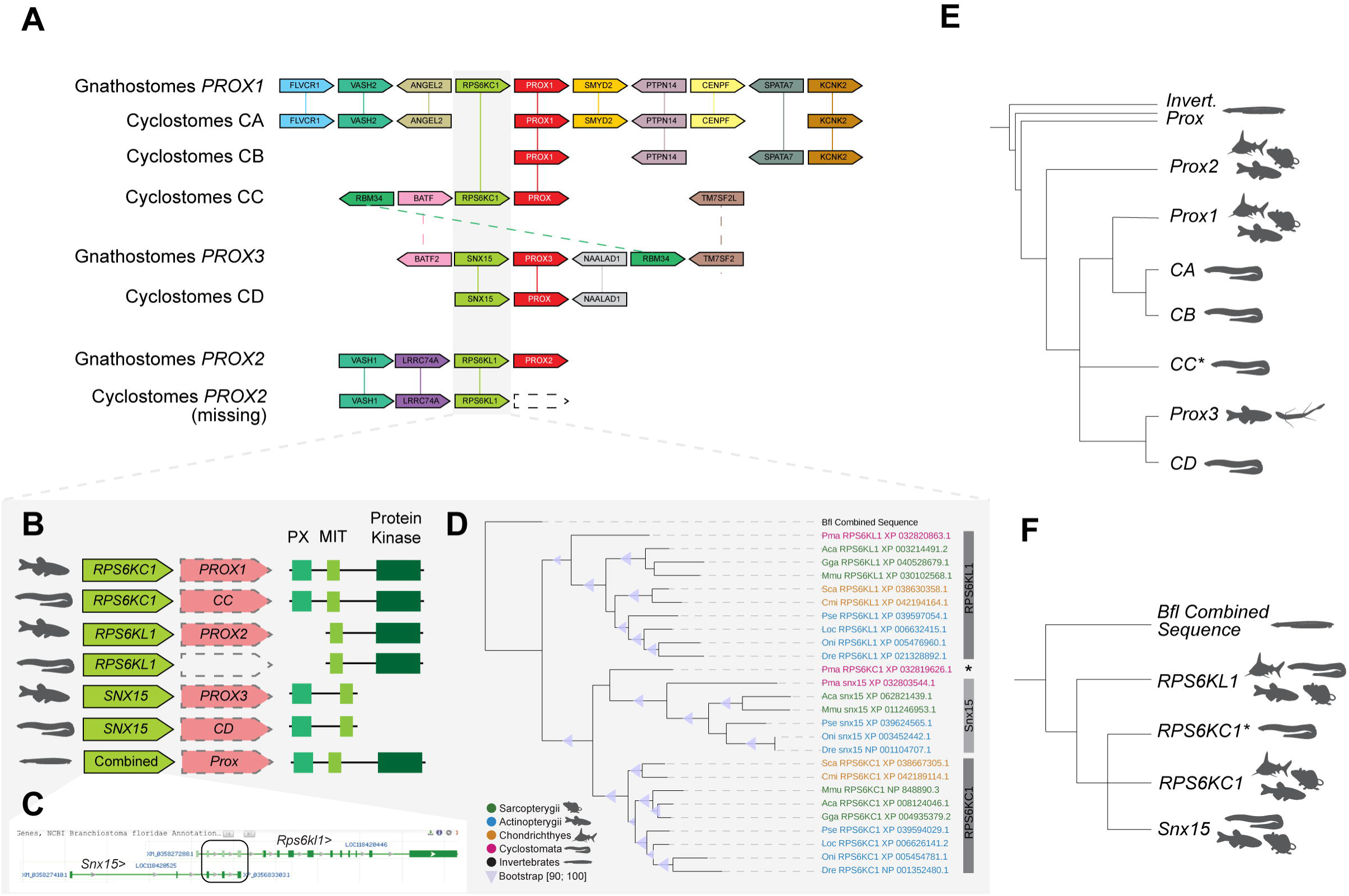
Local synteny indicates the identity of the four cyclostome *Prox* clades. A) Local syntenic conservation between the four cyclostome *Prox* clades and the reconstructed ancestral state of each gnathostome *Prox* clade. Only gene families appearing in more than one clade are shown here. Genes are oriented so that the *Prox* locus is on the forward strand. B) Schematic of the conserved position of the members of the *Rps6kc1/Rps6kl1/Snx15* family upstream of the *Prox* loci and of the conserved protein domains according to Uniprot. C) GenBank annotation for the two adjacent *Rps6kl1/Snx15* loci in *B. floridae* which were combined for this analysis. Box highlights the three exons in common between the two gene predictions. D) Consensus tree of the *Rps6kc1/Rps6kl1/Snx15* family protein sequences based on the Clustal Omega alignment. Nodes with bootstrap value < 50 have been collapsed. Triangles: nodes with [90:100] bootstrap values. Green: sarcopterygian sequences. Blue: actinopterygian sequences. Yellow: chondrichthyan sequences. Magenta: cyclostome sequences. Black: invertebrate/other sequences. Asterisk: cyclostome *Rps6kc1* sequences not retrieved as part of the *Rps6kc1* clade. E) Schematic cladogram of the *Prox* family, with cyclostome clades placed according to conserved synteny. Asterisk: clade with uncertain position F) Schematic cladogram of the *Rps6k* family from the phylogeny in (D). Asterisk: clade with uncertain position

### The three gnathostome Prox subfamilies were present before the divergence of cyclostome and gnathostome lineages

The local syntenic arrangements identified for the gnathostome *Prox* subfamilies (**Figure S2B- D**) were compared to the four cyclostome Prox clades, in order to assign their identity (**Figure 2A**). In Clade B (CB), a second annotated locus positioned in tandem with *Prox* was shown to be homologous to vertebrate *Prox* genes by BLAST search. Upon closer inspection, this gene appears to be a short fragment of the 3’ sequence of the *Prox* gene incorrectly annotated as a separate locus, and was therefore excluded from the successive analysis.

By a comparative approach, several conserved syntenic genes were identified between the *Prox1* subfamily and cyclostome CA and CB (**Figure 2A**). The presence of a *Snx15* locus in CD, whose identity was confirmed by phylogenetic analysis (**Figure 2D**), placed this clade as part of *Prox3* based on local synteny. The presence of a *Prox3* locus in cyclostomes has a substantial implication for the history of this gene, suggesting three independent losses of *Prox3* in vertebrates: in chondrichthyans, in coelacanths, and in tetrapods.

The position of cyclostome CC is less certain. *Petromyzon marinus Rps6kc1* is placed at the base of the vertebrate *Snx15* clade instead of the gnathostome *Rps6kc1* clade (**Figure 2D**), though this placement has weak bootstrap support (<90), suggesting *P. marinus Rps6kc1* could indeed be an ortholog of *Rps6kc1.* If this were the case, it would place CC as part of *Prox1*, and not *Prox3* (**Figure 2A**). However, the rest of the local syntenic landscape close to CC closely matches *Prox3* (**Figure 2A**). This could possibly be due to the retention of features of the ancestral local syntenic organisation in both Prox3 and CC. However, it must be noted that in the phylogenetic analysis CC was retrieved as sister to CD, which has been assigned to the *Prox3* clade. Therefore, the identity of the cyclostome CC clade could not be definitely assigned based on phylogeny and conserved local synteny.

This analysis also uncovered a cyclostome region showing conservation with gnathostomes *Prox2* (**Figure 2A**), containing, among others, genes such as *Rps6kl1*, *Vash1*, and *Lrrc74a*. This is notable, as it suggests that the lack of *Prox2* in cyclostomes is due to a lineage-specific loss, and that the gene was present in the vertebrate ancestor. This is further supported by the presence of both *Prox1* and *Prox3* genes in the cyclostomes, inevitably pushing the split between *Prox2* and the rest of the family before the divergence of the cyclostome and gnathostome lineages.

### Syntenic blocks reconstruction suggests the existence of a lost Prox locus

While local synteny was useful in assigning the identity of the cyclostome *Prox* genes, a chromosome-level synteny block analysis focusing on carefully selected genes was employed to reconstruct the evolutionary scenario of *Prox* gene duplications throughout the history of the vertebrate clade.

We started by considering all the gene families appearing in the analysis in **Figure S2**, and looked for gene families that fulfilled the following criteria: 1) Members of these families are located on the same chromosome as at least two of the *Prox* genes (**Figure S2B-H**), 2) The families include at least three members, and 3) The expansion of the family seems to originate at the base of vertebrates. Gene families following these criteria can be assumed to have originated by the same duplications that gave rise to the *Prox* family, and therefore can help reconstruct the syntenic history of this portion of the genome. We identified six gene families that satisfied our criteria: *Rps6k* (which we already discussed in Figure 2), *Esrr*, *TgX*, *Batf*, *Meis* and *Spred*.

Previous phylogenetic studies have been performed for the *Esrr* (Papadogiannis et al. 2023), *TgX* (S. Liu et al. 2022), *Batf* (Frétaud et al. 2021; Zhu et al. 2019), *Meis* (Irimia et al. 2011) and *Spred* (Motta et al. 2021) families. However, since many key species for synteny reconstruction are not included in the published phylogenies new phylogenetic analyses were run (**Figure S4A-B**). A similar analysis for the *Rps6k* family was already conducted in **Figure 2D**. Sequence identification and phylogenetic analysis were run as previously described. For *Esrr*, *TgX*, *Meis* and *Spred*, the single copies found in invertebrate species were used as an outgroup. For *Batf*, no invertebrate copy was identified, and the closely related *Atf3* family (Frétaud et al. 2021) was used instead. With the exception of *Batf*, three to four monophyletic clades could be identified in all families (**Figure S4A-B**), which further confirmed our selection criteria. However, the cyclostome loci often fell outside such clades (**Figure S4A**) as previously observed for *Prox* (**Figure 1A**). *Batf* phylogenies based on the complete protein failed to identify a monophyletic group corresponding to *Batf2* (data not shown). Since this subfamily is characterized by a long C-terminus tail (Murphy et al. 2013), which could not be aligned to the rest of the sequences, we repeated the analysis using only the highly conserved BZIP domain, present in both *Batf* and *Atf3*. However, this method also failed to retrieve a monophyletic *Batf2* family. It must be noticed that the paraphyletic assemblage contains sequences from all the major gnathostome groups in a single copy (exception made for *D. rerio*, which underwent the additional 3R WGD), which is what would generally be expected to be the case for members of the same subfamily. Alternatively, this paraphyletic assemblage could represent two separate Batf subfamilies, which reciprocally lost their chondrichthyans and osteichthyans members. However, previous studies have identified a monophyletic *Batf2* clade (Zhu et al. 2019). Therefore, this paraphyletic group was considered as *Batf2*, despite the lack of phylogenetic support.

Once sequence identities were confirmed by phylogenetic analysis, their syntenic relation with the identified gnathostome *Prox* loci were investigated (**Figure 3A-D**, **S4B**). Overall, a clear picture emerged, in which members of the different subfamilies are found on the same chromosome as a specific *Prox* gene (**Figure 3A-D**, **S4**). For example, in the *Esrr* family, *Esrrg* is consistently found on the same chromosome with *Prox1*, *Essrb* with *Prox2,* and *Esrra* with *Prox3*.

**Figure 3:**
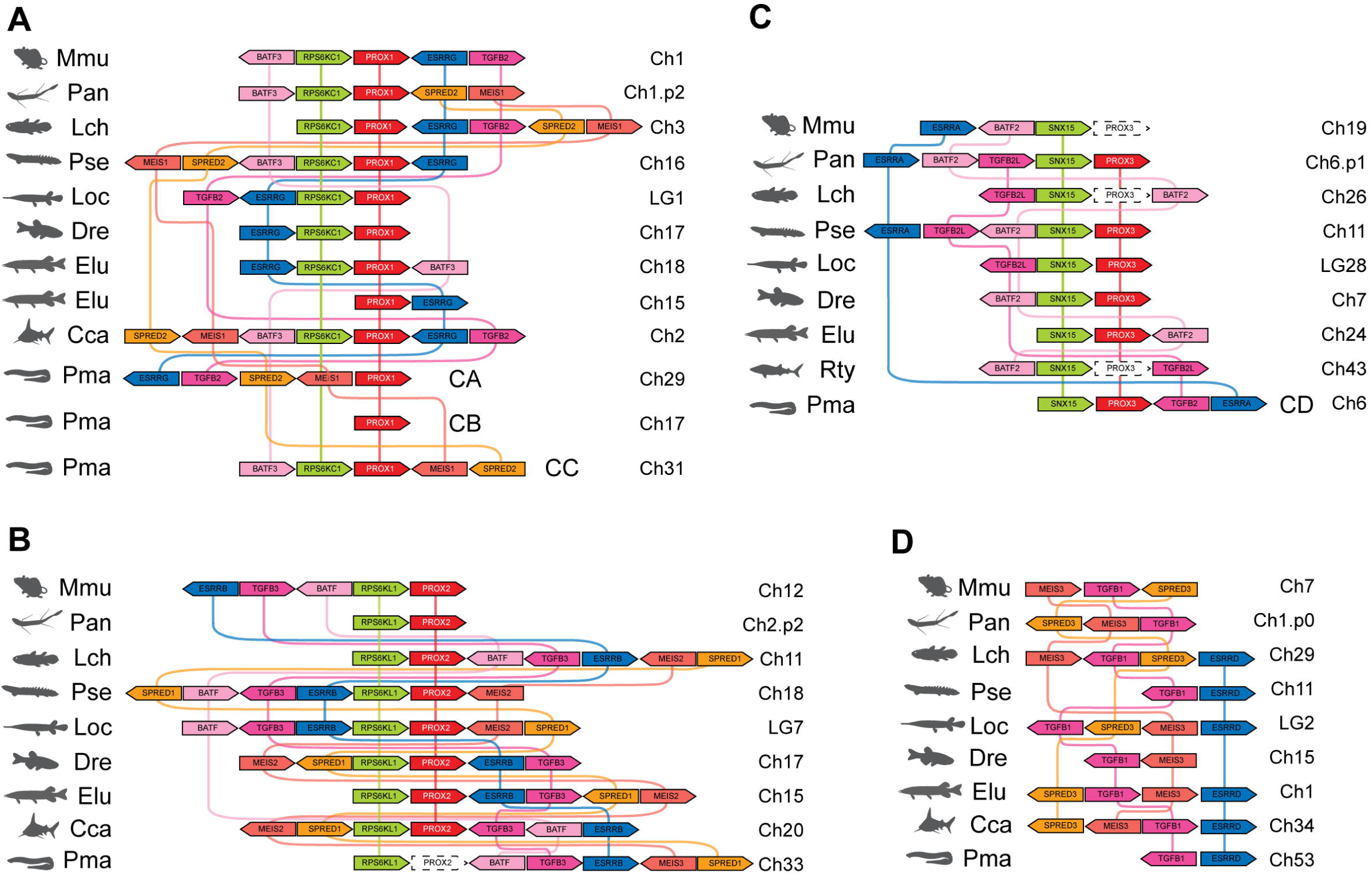
Identification of four conserved syntenic blocks associated with the *Prox* genes. A-D) Position and orientation along the chromosome of the members of the six genes families found associated with the Prox loci: *Spred, Meis, Batf, Rps6k, Esrr, TgX*. Genes are oriented so that the *Prox* locus is on the forward stand. A) Conserved synteny around the *Prox1* locus. B) Conserved synteny around the *Prox2* locus, including the missing *Prox2* in cyclostomes. C) Conserved synteny around the *Prox3* locus, including the missing *Prox3* in chondrichthyans, coelacanth and tetrapods. D) Conserved synteny among members of the *Spred, Meis, TgX* and *Esrr* families in a fourth position, suggesting the presence of a now lost fourth *Prox* locus in all vertebrates.

Through this process, a fourth syntenic block was also identified, marked by the presence of members of the *Esrr*, *TgX*, *Meis* and *Spred* family. Although today no *Prox* locus is found in this region, this likely represents the ancestral syntenic block in which a lost *Prox* gene was located (**Figure 3D**, **S4B**).

### The lost Prox4 syntenic block is common to all major vertebrate clades

Having identified the syntenic blocks corresponding to the *Prox* genes, plus a fourth “lost” one, we used this information to better define the cyclostome Prox genes’ identity.

Cyclostome CA shows conserved synteny with the rest of the *Prox1* clade, and specifically with *TgX2*, whose identity is supported by phylogenetic analysis (**Figure S4A-B**). Other members of the considered families are also found in synteny with this gene, further confirming its identity as part of the *Prox1* subfamily (**Figure 2A**, **S4B, 3A**). Similarly, CD identity as part of *Prox3* is validated by the presence of the *Snx15* locus, as well as members of the *TgX* and *Essr* families. Interestingly, *Spred2* is found in synteny with the *Prox1* and *Rps6kc1* loci in cyclostome CC (**Figure S4B, 3A**), further supporting the identity for this clade as a paralog of *Prox1* and not *Prox3* (**Figure 2A**). This analysis, looking at chromosome-level syntenic conservation and based on the gene families identified in the previous section, found no support for the identity of cyclostome CB as part of the *Prox1* clade. However, in **Figure 2A** we were able to assign CB to *Prox1*, based on more local syntenic conservation, involving genes not found in association with the other *Prox* clades. We can therefore speculate that the *Prox* gene of the CB cyclostome clade has lost its medium- to long-range syntenic conservation as the result of a translocation to a different chromosome. In a similar fashion, almost no hagfish genes were found in these synteny blocks (**Figure 3**, **S4C**), despite the *Prox* loci showing conservation at a more local synteny level, also suggesting extensive chromosomal rearrangements.

Several cyclostome members of the *Esrr*, *TgX*, *Batf*, *Meis* and *Spred* families are found in synteny with what we have identified as the location of the lost cyclostome *Prox2* (**Figure S4B, 3B**), further supporting a gene loss scenario for this locus in cyclostomes.

Interestingly, cyclostome *TgX1* and an unsorted member of the *Essr* family are located on the same chromosome, on which no *Prox* genes are found. As *TgX1* is the member of the *TgX* family found in association with the gnathostome fourth syntenic block (**Figure 3D**), these observations suggest that the same fourth syntenic block identified in Gnathostomes is present in cyclostomes as well.

### Expression of the Prox family is conserved across vertebrates

To investigate the Prox gene functional roles during evolution, we compiled the reported expression domains of the Prox family members across vertebrates (**Figure 4A**, **Table S3**) and several invertebrates (**Figure 4B**, **Table S3**), and analysed the sites of expression shared by at least two *Prox* gene subfamilies. The literature data and publicly available databases used are listed in **Table S3**. Expression was reported when it occurred in the same tissue or organs for at least two Prox genes. When the data were provided at different resolution levels in different organisms, such as for a cell population in one and for the whole organ in another, the more general domain of expression was reported.

**Figure 4:**
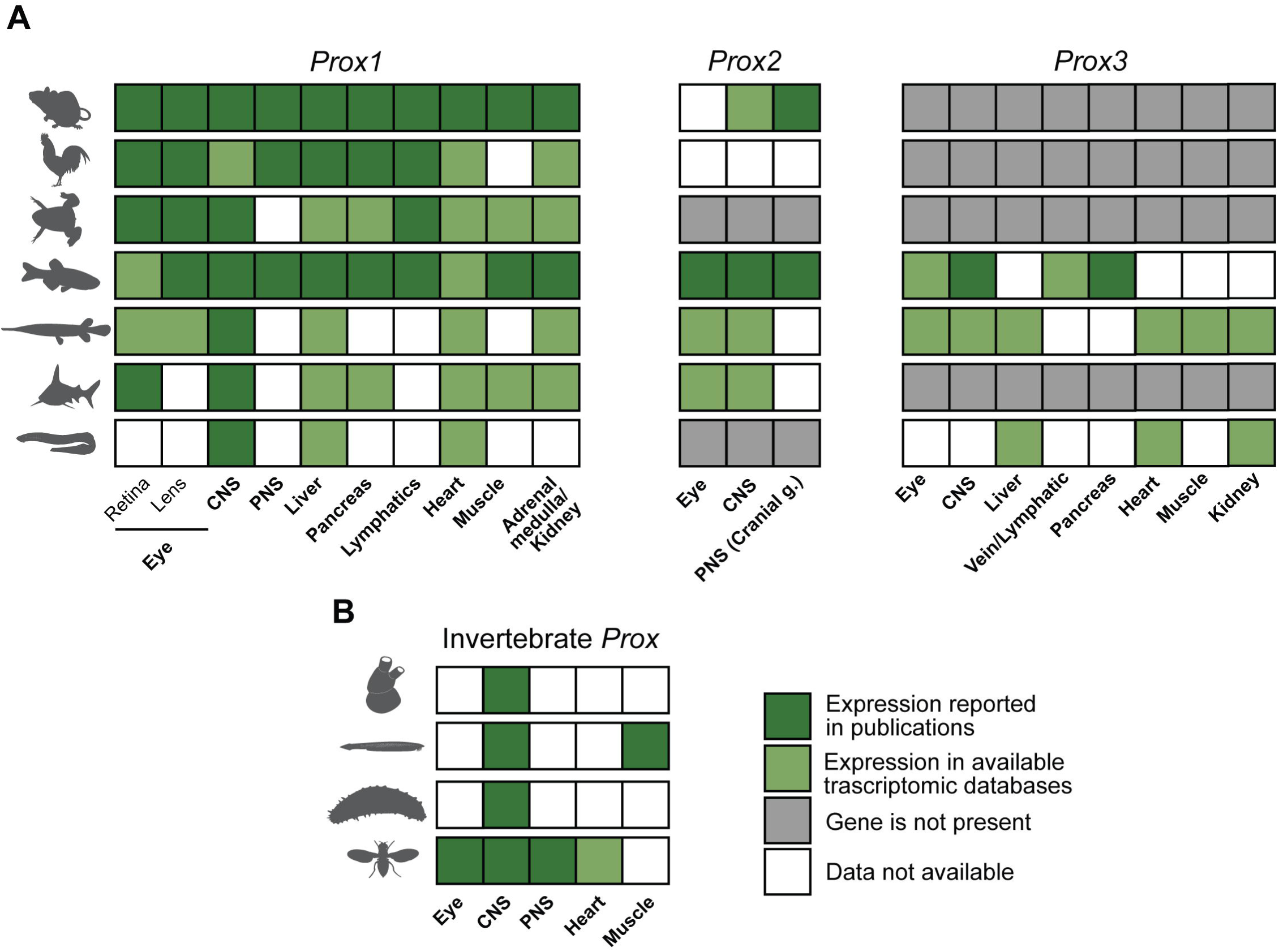
Comparison of the expression of *Prox* genes across vertebrates. A) Conserved expression of members of the *Prox* family in vertebrates. From the top, reported expression in mouse, chicken, Xenopus/newt, zebrafish, spotted gar, lesser spotted catshark and hagfish/lamprey. B) Conserved expression of the *Prox* gene in selected invertebrate species. From the top, sea squirt, lancelet, sea cucumber and fruit fly. Bold: tissues expressing at least two *Prox* subfamilies. Dark green: expression reported in a publication; light green: expression reported in available transcriptomic datasets; grey: gene/organ not present. White: no reported expression/no available data. Relevant references are listed in **Table S3**.

Conserved expression of the *Prox1* genes has been detected in a wide variety of tissues, including neural structures, such as the retina, peripheral nervous system, and central nervous system (CNS). The gene is moreover expressed in the liver and the heart throughout vertebrates, and in the retina, pancreas, and muscles across jawed vertebrates. Expression is also detected in the adrenal medulla of mice, and in the kidney tissue of other species. In bony fishes, where a lymphatic vasculature has been described, *Prox1* is expressed in this tissue (**Figure 4A**). *Prox2* expression seems to be restricted to neuronal structures in all jawed vertebrates, while the gene is missing in cyclostomes. Specifically, expression has been detected in the eye, in the CNS and in the cranial ganglia (**Figure 4A**). As discussed above, *Prox3* is only present in cyclostome and ray-finned fishes. Conserved expression between these groups was described in the liver, heart, and kidney, while the gene is also expressed in the vasculature, pancreas, neural structures and muscles of ray-finned fishes (**Figure 4A**). The expression of *Prox/prospero* genes was also reported for the available invertebrate species, where a conserved expression in the CNS was detected (**Figure 4B**).

Overall, a high level of redundancy in expression domains can be seen between *Prox1* and *Prox3*, while *Prox2* seems to be restricted to neuronal structures, in a pattern more similar to that on invertebrates.

### Possible models of Prox family evolution

Based on the phylogenetic analyses and conserved synteny reconstructions in cyclostomes, we reconstructed two alternative models of evolution of the *Prox* family. The first model (**Figure 5A**) assumes that the basal split of the vertebrate *Prox* family is the result of a gene duplication, followed by the 1R WGD. In this model, four copies of *Prox* were present in the lineage leading to vertebrates, and *Prox4* was lost before the split between cyclostomes and gnathostomes. In gnathostomes, *Prox3* was subsequently lost in several lineages (chondrichthyans, coelacanths, tetrapods, and one of the copies from the teleost WGD). In cyclostomes, *Prox* family members from both the *Prox1* and *Prox3* clades can be found, while *Prox2* has been lost in this lineage. The three copies of *Prox1* found in cyclostomes are due to the genome triplication that happened in the ancestors of this group. Accordingly, this model also postulates the loss of two additional cyclostome *Prox3* genes, following the duplication.

**Figure 5:**
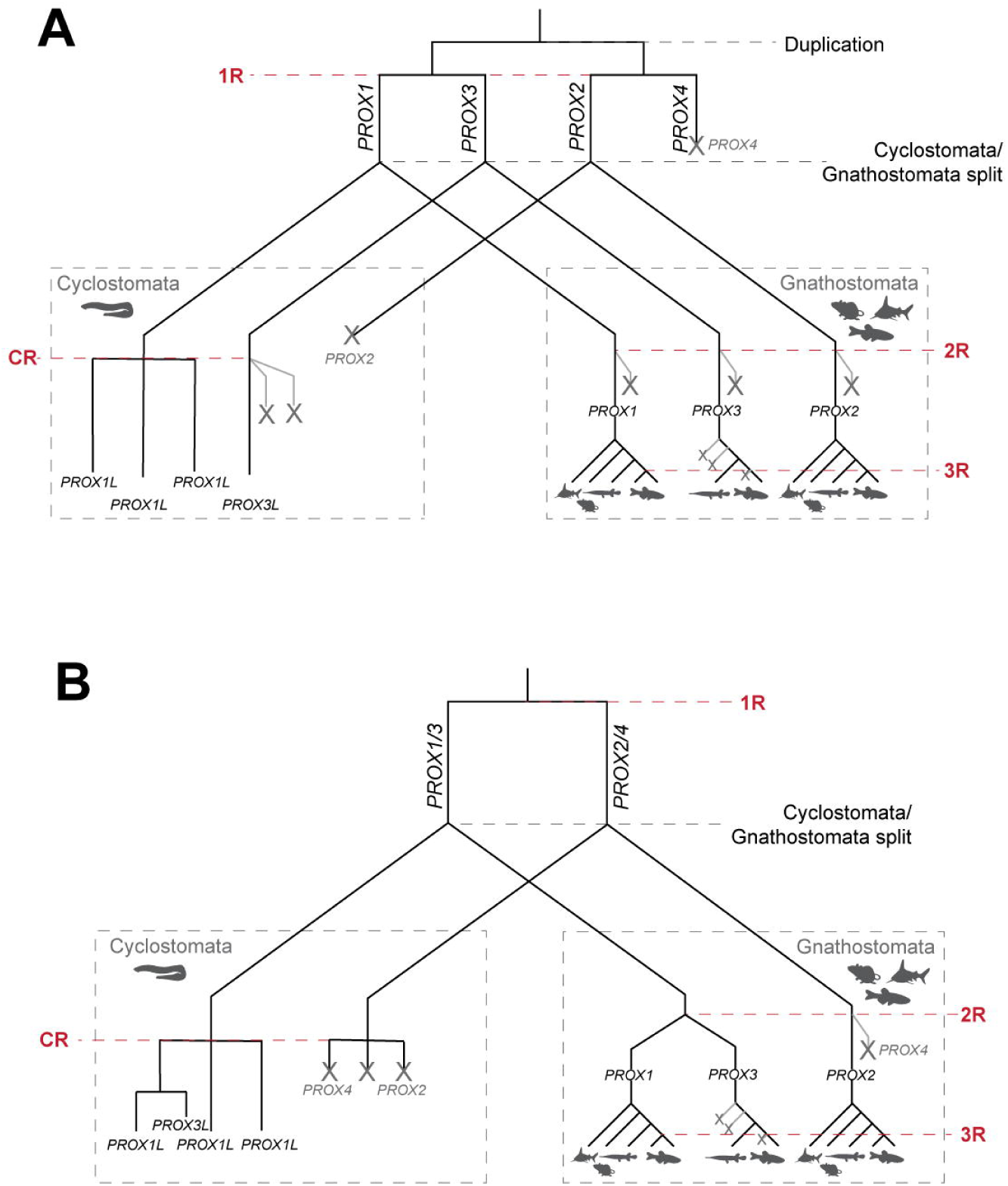
Alternative models of the *Prox* family evolution. A) At the base of the vertebrate *Prox* family is a local duplication. Subsequently, the 1R WGD give raise to *Prox1* and *Prox3* on one hand, and *Prox2* and a nowadays lost *Prox4* on the other. In this model, *Prox4* would have been present the ancestors of modern vertebrates, and subsequently lost. B) The duplication at the base of the vertebrate *Prox* family is the result of the 1R WGD, and only two Prox genes were present in the ancestors of modern vertebrates. In this model, the genes identified by the same names in gnathostome and cyclostome are not orthologues, as they arise from independent duplication events in these lineages.

An alternative scenario (**Figure 5B**) postulates that two *Prox* genes, the ancestor of *Prox1* and *Prox3* (here called *Prox1/3*) and that of *Prox2* and *Prox4* (*Prox2/4*) emerged from the 1R WGD and were present at the split between cyclostomes and gnathostomes. In this scenario, the four gnathostome genes emerged from the 2R WGD, with *Prox4* then being lost, while the four cyclostome genes are the results of the genome triplication in cyclostome and an additional gene duplication. The *Prox2* and *Prox4* cyclostome genes are similarly the result of the triplication of the *Prox2/4* gene, and were lost afterwards. It is important to note that in this scenario, the current nomenclature would be incorrect, as for example, cyclostome *Prox3* (CD) would not be the ortholog of gnathostome *Prox3,* but more closely related to the cyclostome *Prox1* sequences.

## Discussion

### The evolutionary history of the Prox family throughout bilaterians

While homeobox genes are found in most Eukaryotes, genes of the PROS class, which includes both *prospero* and *Prox*, are only found in Bilateria (Holland 2013). Homeobox genes of the PROS class are characterised by the presence of a highly divergent homeodomain, lacking basic residues at the N-terminus (Bürglin and Affolter 2016), and present the PROSPERO domain at the C-terminus of the protein sequence (Bürglin 1994) .

Within bilaterians, members of the PROS class have been identified in arthropods (Stollewerk 2016), such as the fruit fly *Drosophila melanogaster* (Doe et al. 1991), the water flea *Daphnia magna* (Ungerer et al. 2011), and the spider *Cupiennius salei* (Weller and Tautz 2003), as well as other ecdysozoans such as the nematode *Caenorhabditis elegans* (Bürglin 1994). PROS genes are also present in lophotrochozoans, as shown by the presence of a *prospero* gene in the anellids *Malacoceros fuliginosus* and *Platynereis dumerilii* (Kerner et al. 2009; Kumar et al. 2020) and the cephalopod *Doryteuthis pealeii* (Koenig et al. 2016).

In deuterostomes, one copy of the *Prox/prospero* gene is found in non-vertebrate species, such as the acorn worm *Saccoglossus kowalevskii* (Freeman et al. 2008), and in lancelet (Zawisza-Álvarez et al. 2020). In this work, we found that two *Prox* genes are present in tandem in the purple sea urchin *Strongylocentrotus purpuratus.* However, the phylogenetic analysis (**Figure 1A**) suggests this is a very recent lineage-specific event.

*prospero* genes are also found in Xenacoelomorpha (Brauchle et al. 2018), which is considered the sister group of all bilaterians (Rouse et al. 2016; Cannon et al. 2016), further supporting the idea of an origin of PROS genes in the lineage leading to bilaterians.

### Reconstruction of the Prox family distribution and history in vertebrates

Within vertebrates, an expansion of the Prox family can be observed. In this study, we confirmed that three *Prox* subfamilies are present in gnathostomes, named *Prox1*, *Prox2,* and *Prox3* (**Figure 1A-B**). The latter has a somewhat limited phylogenetic distribution, being found only in actinopterygians and a single sarcopterygian species, the lungfish *Protopterus annectens.* Prox1 and Prox3 are sister clades to each other, with Prox2 lying outside of them. Four Prox genes can also be identified in cyclostomes. Local synteny analysis suggests these genes are closely related to the *Prox1* and *Prox3* clade, while the *Prox2* gene seems to have been lost in cyclostomes (**Figure 2A-B**).

Chromosome-level synteny is a powerful tool to employ together with phylogeny to establish the evolutionary history and relationship of gene families and subfamilies. In our study, we characterised the co-occurrence of six gene families, Rps6kc, Esrr, Tgfb, Batf, Meis and Spred together with the different Prox genes. Specific members of these families are consistently found in association with a Prox subfamily: for example, *Rps6kc1*, *Essrg, TgX2*, *Batf3, Meis1* and *Spred2* are often located on the same chromosome as *Prox1*. Based on this chromosome-level synteny approach, the presence of a fourth syntenic block, which we reason was the site of a fourth *Prox* gene now lost, was identified in both gnathostomes and cyclostomes (**Figure 3D**). In addition, a previous investigation of the muscarinic acetylcholine receptors family, whose members are located in synteny with *Prox* genes, identified other gene families colocalising with the reconstructed *Prox* synteny blocks, such as EDH, LTBP and PRKD (Pedersen et al. 2018). Remarkably, these families also include members localising on the same chromosome as the reconstructed *Prox4* locus, strengthening our observations.

### Uncovering the evolutionary history of the Prox gene family in vertebrates: insights from syntenic loci

Based on the data and observation above, we proposed two possible scenarios of the evolution of the Prox family in vertebrates, based on the integration of phylogeny and synteny analysis.

In the first scenario (**Figure 5A**), we hypothesize that the expansion of the Prox family in vertebrates is driven at first by a duplication event, followed by the 1R WGD. This scenario postulates the presence of four *Prox* genes in the lineage leading to vertebrates, with Prox4 being lost sometimes before the split between cyclostomes and gnathostomes. In this scenario, three of the cyclostome *Prox* genes are paralogs to gnathostome *Prox1*, one is an ortholog of gnathostome *Prox3*, while *Prox2* was present at the base of cyclostomes and later lost.

A second possible scenario (**Figure 5B**) does not rely on any additional duplication and instead postulates that only two Prox genes, corresponding to the ancestors of *Prox1/Prox3* and *Prox2/Prox4*, were present at the base of vertebrates. In this scenario, gnathostome *Prox1* and *Prox3* do not have one-to-one orthologues in cyclostomes, as the *Prox* genes present in the latter would have emerged from the cyclostome-specific genome triplication event.

In order to evaluate which scenario is more likely, local and chromosome-level synteny should be considered, looking at the co-occurrences of genes around *Prox1* and *Prox3*. Such genes are of key interest, as in the second scenario, any similarity shared only between gnathostome *Prox1* or *Prox3* and a cyclostome *Prox* gene must be explained by either parallel retention of ancestral characters or parallel evolution of new ones. In other words, in the second scenario we would expect gnathostome *Prox1* to share more syntenic similarities with gnathostome *Prox3* than with cyclostome *Prox* genes.

Looking at the local synteny (**Figure 2A**), the parallel loss of several loci, including *Flvcr1*, *Vash2*, *Angel2*, *Smyd2*, and *Naalad1*, would be needed to support the second scenario. However, this possibility cannot be excluded, as even assuming the first scenario, several of these genes must have been lost in cyclostome CB and CC *Prox1* genes (**Figure 2A**). Similarly, no definitive proof in support of one model or the other can be drawn from the chromosome- level synteny blocks, as the attribution of the cyclostome genes’ identity has very low support for most gene families.

The strongest support for one model over the other comes from the distribution of the *Rps6K* family (**Figure 2B-D**). As stated before, members of this family are found in association with *Prox* genes throughout vertebrates. Namely, *Rps6kc1* and *Snx15* are found in synteny with Gnathostome *Prox1* and *Prox3*, respectively, as well as two of the cyclostome *Prox* genes. This observation supports the first model over the second, as it implies that both *Prox1* and *Prox3*, associated with *Rps6kc1* and *Snx15,* were present at the split between cyclostome and gnathostome. Otherwise, a scenario in which *Snx15* evolved twice from *Rps6kc1* by the loss of the protein kinase domain (**Figure 2B**) should be proposed, which contradicts the phylogenetic results (**Figure 2D**). For these reasons, we propose that the first model of the *Prox* family evolution, which postulates the presence of three Prox genes at the split of cyclostome and gnathostome, is better supported.

### Prox genes and the 3R WGD in teleosts: implication for current research

During their evolutionary history, teleost fishes underwent a lineage-specific WGD event, called 3R. As a result of this, many genes are found in two copies, or ohnologues, in teleosts. Two copies of *Prox1*, called *prox1a* and *prox1b*, are found in several teleost species (**Figure 1A**, **S2B**), but crucially, only one gene, *prox1a*, is present in the model organism *Danio rerio*. Similarly, although *Prox2* is present in two copies in some teleosts, only one *prox2* gene is found in *Danio rerio* (**Figure 1A**, **S2C**). In this study, we could only identify one *Prox3* gene in teleosts, suggesting the second copy emerging from 3R was almost immediately lost (**Figure 1A**, **S2D**). This single *prox3* gene has in the past being identified as the second *Prox1* ohnologues, *prox1b* (Giacco et al. 2010), but both phylogeny and synteny clearly identify this gene as *prox3*. This observation is particularly relevant as *prox1b/prox3* has been a subject of some interest in the field of lymphatic vascular research (Tao et al. 2011; van Impel et al. 2014; Koltowska et al. 2015), as *Prox1/prox1a* has a well-documented role in the development and maintenance of lymphatic vessels (Wigle and Oliver 1999; Nicenboim et al. 2015; Koltowska et al. 2015). Indeed, there is evidence that *prox3* is able to partially compensate for the loss of *prox1a* in lymphatic vascular development (Grimm et al. 2023). The ability of *prox3* to compensate for *prox1a* loss was assumed to stem from the high redundancy between ohnologues from the 3R WGD. Instead, our study shows that the divergence between the two genes potentially dates back to the base of the vertebrate clade (**Figure 4**), suggesting a wider role of the *Prox* family in lymphatic development across vertebrates.

### Functional evolution of Prox genes

Having established the phylogenetic relationship and evolutionary history of the members of the vertebrate Prox family, we leveraged this knowledge in combination with gene expression profiles to shed new light on the evolution of *Prox* gene functional roles.

To investigate this, the reported expression domains of the Prox family members across vertebrates (**Figure 4A**, **Table S3**) and several invertebrates (**Figure 4B**, **Table S3**) were compiled. As expected, *Prox1* members show a wide expression profile described in the literature (listed in **Table S3**), while *Prox2* genes are expressed in the structures of the nervous system, in agreement with their described role in the vagal neurons and enteric ganglia associated with oesophageal mobility (referenced in **Table S3**). Despite having been lost in several lineages, *Prox3* is documented in the transcriptomic datasets of *Danio rerio, Lepisosteus oculatus*, and *Eptatretus burgeri* as having a wide expression pattern in several organs, not dissimilar to *Prox1*.

From these data, we can reconstruct that expression in neural structures (such as eyes, CNS and PNS) appears to be ancestral in *Prox* genes, as such expression is detected across all investigated subfamilies as well as in invertebrate species. *Prox2* expression is also restricted to neuronal structures, suggesting this might have been the function of the ancestor of vertebrate *Prox* genes. However, both *Prox1* and *Prox3* show expanded and often overlapping patterns of expression that include liver, pancreas, venous/lymphatic vasculature, heart, muscle, and kidney structures. This suggests that before the whole-genome duplication at the base of vertebrates, the ancestor of *Prox1* and *Prox3* had already expanded its function to these organs. Overall, when combined with the reconstructed evolutionary history of the *Prox* family (**Figure 5A**), these data suggest a scenario in which the ancestral nervous system-related expression of *Prox* genes has been maintained in all vertebrate *Prox* subfamilies (**Figure 4A-B**, **Graphical abstract**). Meanwhile, an expansion of the expression domains has happened before the duplication of *Prox1* and *Prox3*, leading to a high degree of redundancy between these genes, which might explain the repeated loss of *Prox3* in gnathostome lineages (**Figure 4A-B**, **Graphical abstract**).

## Conclusions

In this study we have produced a full characterisation of the evolutionary history of the *Prox* gene family, reconstructing the presence of four *Prox* genes at the base of vertebrate, one of which is now lost in all extant lineages. Moreover, we leveraged published expression data to provide a scenario for the evolution of *Prox* gene functions in vertebrates, reconstructing an expansion of the expression domains in the ancestor of the *Prox1* and *Prox3* clades. This work represents the first extensive description of the *Prox* family evolutionary history in vertebrates, clarifying past misidentifications of its members and laying the foundations for future studies on the function of these genes.

## Methods

### Phylogenetic analysis

Protein sequences were retrieved from GenBank genome databases for invertebrate and vertebrate species, and Ensembl database for hagfish (*Eptatretus burgeri*). All species used, genome assembly identifiers, and accession numbers are listed in **Table S1** for the *Prox* phylogeny and in **Table S2** for the other gene family phylogenies.

Sequence alignment was performed using Seaview 5.0.5 (Gouy et al. 2010; Galtier et al. 1996). with either Clustal Omega (Sievers et al. 2011) or MUSCLE (Edgar 2004). Long, unaligned sequences were trimmed. Phylogenetic analyses were conducted locally using IQ-TREE2 (Minh et al. 2020) with ModelFinder (Kalyaanamoorthy et al. 2017) and SH-aLRT test (n=1000). Bootstrapping was performed using UFBoot (Hoang et al. 2018) (n=1000). Consensus tree visualisation was carried out using iTOL (Letunic and Bork 2021). Nodes with bootstrap values < 50 were collapsed into polytomies, while nodes with bootstrap values between 90 and 100 were considered fully supported.

### Conserved local synteny analysis

Genomic regions containing up to ten neighbouring genes upstream and downstream of each Prox gene sequence were retrieved using the EzGeneSynteny package, described below. BLAST against GenBank and Ensembl was used to confirm the collected synteny and assign gene identity when annotation was not complete. Only gene families sharing the loci with at least two *Prox* clades were further considered in this analysis.

Consensus synteny for a subset of species (**Figure 2A**) was determined based on principles of parsimony from the blocks in **Figure S2**. Synteny shared with the invertebrate outgroup, such as the placement of *Naalad1* downstream of *Prox3*, was noted in the figure but not considered in assigning the clade identity. Illustrated with Adobe Illustrator 2026 (Adobe Inc.), based on the output of *EzGeneSynteny*.

### Reconstruction of the syntenic blocks

Gene identity within the considered gene families was assigned by phylogenetic analysis (see above). The position of the loci on the chromosome was checked using GenBank. Ancestral Prox synteny blocks were predicted based on principles of parsimony. Illustrated with Adobe Illustrator 2026 (Adobe Inc.).

### Generation of EzGeneSynteny package

The EzGeneSynteny package is a tool developed for simple and rapid visualization of gene order across multiple genomes through a simple command-line interface that requires no local genome files (https://github.com/jakeleyhr/EZgenesynteny). The program accepts user- defined sets of orthologous or homologous genes and retrieves the corresponding genomic regions and annotations from NCBI GenBank via the Entrez API. For each input gene, *EZgenesynteny* extracts the surrounding genomic context, parses all annotated coding sequence (CDS) features, and orders the user-specified number of upstream and downstream genes according to their chromosomal coordinates. *EZgenesynteny* outputs both a CSV file and a synteny plot to easily compare the conservation of gene order across species.

### Transcriptome data mining

Reported expression of Prox genes was investigated in a selection of vertebrate and invertebrate species: mouse (*Mus musculus*), chicken (*Gallus gallus*), African clawed frog (*Xenopus laevis*), western clawed frog (*Xenopus tropicalis*), newt (*Pleurodeles waltl*), zebrafish (*Danio rerio*), spotted gar (*Lepisosteus oculatus*), lesser spotted catshark (*Scyliorhinus canicula*), river lamprey (*Lampetra fluviatilis*), sea lamprey (*Petromyzon marinus*), inshore hagfish (*Eptatretus burgeri*), seasquirt (*Ciona robusta),* amphioxus *(Branchiostoma lanceolatum),* sea cucumber *(Holothuria glaberrima),* and fruit fly *(Drosophila melanogaster).* First, a literature search was conducted to identify published expression data for Prox genes (results listed in **Table S3**). This data were then integrated with information from publicly available gene expression databases (listed in **Table S3**). Expression reported only in genome-wide databases and only for one Prox gene were not included.

## Acknowledgments

We thank Professor Melanie Debiais-Thibaud for providing access to the *S. canicula* transcriptomic data.

## Author Contributions

V. P., T. H. and K.K. participated in the design of the study. V.P. performed the analyses and database searches. J. L. developed the EzGeneSynteny package. V. P. and T. H. performed the data interpretation. V. P. and T. H. wrote the manuscript, with input from the other co-authors.

## Data availability statement

The EzGeneSynteny package is publicly available at https://github.com/jakeleyhr/EZgenesynteny.

## Literature cited

1. Allendorf, FW, and GH Thorgaard. 1984. “Tetraploidy and the Evolution of Salmonid Fishes.” In Evolutionary Genetics of Fishes. Plenum Publishing Corporation.

2. Becker, Jürgen, Baigang Wang, Helena Pavlakovic, Kerstin Buttler, and Jörg Wilting. 2010. “Homeobox Transcription Factor Prox1 in Sympathetic Ganglia of Vertebrate Embryos: Correlation With Human Stage 4s Neuroblastoma.” Pediatric Research 68 (2): 112–17. 10.1203/PDR.0b013e3181e5bc0f.

3. Brauchle, Michael, Adem Bilican, Claudia Eyer, et al. 2018. “Xenacoelomorpha Survey Reveals That All 11 Animal Homeobox Gene Classes Were Present in the First Bilaterians.” Genome Biology and Evolution 10 (9): 2205–17. 10.1093/gbe/evy170.

4. Bürglin, Thomas R. 1994. “A Caenorhabditis Elegans Prosperohomologue Defines a Novel Domain.” Trends in Biochemical Sciences 19 (2): 70–71. 10.1016/0968-0004(94)90035-3.

5. Bürglin, Thomas R., and Markus Affolter. 2016. “Homeodomain Proteins: An Update.” Chromosoma 125 (3): 497–521. 10.1007/s00412-015-0543-8.

6. Burke, Zoë, and Guillermo Oliver. 2002. “*Prox1* Is an Early Specific Marker for the Developing Liver and Pancreas in the Mammalian Foregut Endoderm.” Mechanisms of Development 118 (1): 147–55. 10.1016/S0925-4773(02)00240-X.

7. Cannon, Johanna Taylor, Bruno Cossermelli Vellutini, Julian Smith, Fredrik Ronquist, Ulf Jondelius, and Andreas Hejnol. 2016. “Xenacoelomorpha Is the Sister Group to Nephrozoa.” Nature 530 (7588): 89–93. 10.1038/nature16520.

8. Cardoso, João C. R., Christina A. Bergqvist, and Dan Larhammar. 2020. “Corticotropin-Releasing Hormone (CRH) Gene Family Duplications in Lampreys Correlate With Two Early Vertebrate Genome Doublings.” Frontiers in Neuroscience 14 (July). 10.3389/fnins.2020.00672.

9. Chen, Zelin, Yoshihiro Omori, Sergey Koren, et al. 2019. “De Novo Assembly of the Goldfish (Carassius Auratus) Genome and the Evolution of Genes after Whole-Genome Duplication.” Science Advances 5 (6): eaav0547. 10.1126/sciadv.aav0547.

10. Doe, Chris Q., Quynh Chu-LaGraff, Dorothy M. Wright, and Matthew P. Scott. 1991. “The Prospero Gene Specifies Cell Fates in the Drosophila Central Nervous System.” Cell 65 (3): 451–64. 10.1016/0092-8674(91)90463-9.

11. Dyer, Michael A., Frederick J. Livesey, Constance L. Cepko, and Guillermo Oliver. 2003. “Prox1 Function Controls Progenitor Cell Proliferation and Horizontal Cell Genesis in the Mammalian Retina.” Nature Genetics 34 (1): 53–58. 10.1038/ng1144.

12. Edgar, Robert C. 2004. “MUSCLE: Multiple Sequence Alignment with High Accuracy and High Throughput.” Nucleic Acids Research 32 (5): 1792–97. 10.1093/nar/gkh340.

13. Elsir, Tamador, Anja Smits, Mikael S. Lindström, and Monica Nistér. 2012. “Transcription Factor PROX1: Its Role in Development and Cancer.” Cancer and Metastasis Reviews 31 (3–4): 793–805. 10.1007/s10555-012-9390-8.

14. Ford, Caitlin, Carmen de Sena-Tomás, Tint Tha Ra Wun, et al. 2025. “Nkx2.7 Is a Conserved Regulator of Craniofacial Development.” Nature Communications 16 (1): 3802. 10.1038/s41467-025-58821-3.

15. Freeman, R. M., M. Wu, M. M. Cordonnier-Pratt, et al. 2008. “cDNA Sequences for Transcription Factors and Signaling Proteins of the Hemichordate Saccoglossus Kowalevskii: Efficacy of the Expressed Sequence Tag (EST) Approach for Evolutionary and Developmental Studies of a New Organism.” The Biological Bulletin 214 (3): 284–302. 10.2307/25470670.

16. Frétaud, Maxence, Nam Do Khoa, Armel Houel, Aurélie Lunazzi, Pierre Boudinot, and Christelle Langevin. 2021. “New Reporter Zebrafish Line Unveils Heterogeneity among Lymphatic Endothelial Cells during Development.” Developmental Dynamics 250 (5): 701–16. 10.1002/dvdy.286.

17. Galtier, N., M. Gouy, and C. Gautier. 1996. “SEAVIEW and PHYLO_WIN: Two Graphic Tools for Sequence Alignment and Molecular Phylogeny.” Bioinformatics 12 (6): 543–48. 10.1093/bioinformatics/12.6.543.

18. Garcia-Concejo, Adrian, and Dan Larhammar. 2021. “Protein Kinase C Family Evolution in Jawed Vertebrates.” Developmental Biology 479 (November): 77–90. 10.1016/j.ydbio.2021.07.013.

19. Giacco, Luca Del, Anna Pistocchi, and Anna Ghilardi. 2010. “Prox1b Activity Is Essential in Zebrafish Lymphangiogenesis.” PLOS ONE 5 (10): e13170. 10.1371/journal.pone.0013170.

20. Gouy, Manolo, Stéphane Guindon, and Olivier Gascuel. 2010. “SeaView Version 4: A Multiplatform Graphical User Interface for Sequence Alignment and Phylogenetic Tree Building.” Molecular Biology and Evolution 27 (2): 221–24. 10.1093/molbev/msp259.

21. Grimm, Lin, Elizabeth Mason, Hujun Yu, et al. 2023. “Single-Cell Analysis of Lymphatic Endothelial Cell Fate Specification and Differentiation during Zebrafish Development.” The EMBO Journal 42 (11): e112590. 10.15252/embj.2022112590.

22. Hoang, Diep Thi, Olga Chernomor, Arndt von Haeseler, Bui Quang Minh, and Le Sy Vinh. 2018. “UFBoot2: Improving the Ultrafast Bootstrap Approximation.” Molecular Biology and Evolution 35 (2): 518–22. 10.1093/molbev/msx281.

23. Holland, Peter W. H. 2013. “Evolution of Homeobox Genes.” WIREs Developmental Biology 2 (1): 31–45. 10.1002/wdev.78.

24. Holzmann, Julia, Melanie Hennchen, and Hermann Rohrer. 2015. “Prox1 Identifies Proliferating Neuroblasts and Nascent Neurons during Neurogenesis in Sympathetic Ganglia.” Developmental Neurobiology 75 (12): 1352–67. 10.1002/dneu.22289.

25. Impel, Andreas van, Zhonghua Zhao, Dorien M. A. Hermkens, et al. 2014. “Divergence of Zebrafish and Mouse Lymphatic Cell Fate Specification Pathways.” Development 141 (6): 1228–38. 10.1242/dev.105031.

26. Irimia, Manuel, Ignacio Maeso, Demián Burguera, et al. 2011. “Contrasting 5’ and 3’ Evolutionary Histories and Frequent Evolutionary Convergence in Meis/Hth Gene Structures.” Genome Biology and Evolution 3 (January): 551–64. 10.1093/gbe/evr056.

27. Kaltezioti, Valeria, Georgia Kouroupi, Maria Oikonomaki, et al. 2010. “Prox1 Regulates the Notch1-Mediated Inhibition of Neurogenesis.” PLoS Biology 8 (12): e1000565. 10.1371/journal.pbio.1000565.

28. Kalyaanamoorthy, Subha, Bui Quang Minh, Thomas K. F. Wong, Arndt von Haeseler, and Lars S. Jermiin. 2017. “ModelFinder: Fast Model Selection for Accurate Phylogenetic Estimates.” Nature Methods 14 (6): 587–89. 10.1038/nmeth.4285.

29. Kerner, Pierre, Elena Simionato, Martine Le Gouar, and Michel Vervoort. 2009. “Orthologs of Key Vertebrate Neural Genes Are Expressed during Neurogenesis in the Annelid Platynereis Dumerilii.” Evolution & Development 11 (5): 513–24. 10.1111/j.1525-142X.2009.00359.x.

30. Kivelä, Riikka, Ida Salmela, Yen Hoang Nguyen, et al. 2016. “The Transcription Factor Prox1 Is Essential for Satellite Cell Differentiation and Muscle Fibre-Type Regulation.” Nature Communications 7 (1): 13124. 10.1038/ncomms13124.

31. Koenig, Kristen M., Peter Sun, Eli Meyer, and Jeffrey M. Gross. 2016. “Eye Development and Photoreceptor Differentiation in the Cephalopod *Doryteuthis Pealeii*.” *Development*, January 1, dev.134254. 10.1242/dev.134254.

32. Koltowska, Katarzyna, Anne Karine Lagendijk, Cathy Pichol-Thievend, et al. 2015. “Vegfc Regulates Bipotential Precursor Division and Prox1 Expression to Promote Lymphatic Identity in Zebrafish.” Cell Reports 13 (9): 1828–41. 10.1016/j.celrep.2015.10.055.

33. Kumar, Suman, Sharat Chandra Tumu, Conrad Helm, and Harald Hausen. 2020. “The Development of Early Pioneer Neurons in the Annelid Malacoceros Fuliginosus.” BMC Evolutionary Biology 20 (1): 117. 10.1186/s12862-020-01680-x.

34. Kuraku, S., A. Meyer, and S. Kuratani. 2008. “Timing of Genome Duplications Relative to the Origin of the Vertebrates: Did Cyclostomes Diverge before or After?” Molecular Biology and Evolution 26 (1): 47–59. 10.1093/molbev/msn222.

35. Lanoizelet, Maxence, Léo Michel, Ronan Lagadec, et al. 2024. “Analysis of a Shark Reveals Ancient, Wnt- Dependent, Habenular Asymmetries in Vertebrates.” Nature Communications 15 (1): 10194. 10.1038/s41467-024-54042-2.

36. Lavado, Alfonso, and Guillermo Oliver. 2007. “*Prox1* Expression Patterns in the Developing and Adult Murine Brain.” Developmental Dynamics 236 (2): 518–24. 10.1002/dvdy.21024.

37. Letunic, Ivica, and Peer Bork. 2021. “Interactive Tree Of Life (iTOL) v5: An Online Tool for Phylogenetic Tree Display and Annotation.” Nucleic Acids Research 49 (W1): W293–96. 10.1093/nar/gkab301.

38. Li, Ruihan, Xiaoai Wang, Chao Bian, et al. 2021. “Whole-Genome Sequencing of Sinocyclocheilus Maitianheensis Reveals Phylogenetic Evolution and Immunological Variances in Various Sinocyclocheilus Fishes.” Frontiers in Genetics 12 (October). 10.3389/fgene.2021.736500.

39. Liu, Siqi, Junfu Guo, Xianda Cheng, et al. 2022. “Molecular Evolution of Transforming Growth Factor-β (TGF-β) Gene Family and the Functional Characterization of Lamprey TGF-Β2.” Frontiers in Immunology 13 (March). 10.3389/fimmu.2022.836226.

40. Liu, Xin, Honghui Zeng, Cheng Wang, et al. 2022. “Improved Genome Assembly of Chinese Sucker (Myxocyprinus Asiaticus) Provides Insights into the Identification and Characterization of Pharyngeal Teeth Related Maker Genes in Cyprinoidei.” Water Biology and Security 1 (3): 100049. 10.1016/j.watbs.2022.100049.

41. Lowenstein, Elijah D., Pierre-Louis Ruffault, Aristotelis Misios, et al. 2023. “Prox2 and Runx3 Vagal Sensory Neurons Regulate Esophageal Motility.” Neuron 111 (14): 2184–2200.e7. 10.1016/j.neuron.2023.04.025.

42. Minh, Bui Quang, Heiko A. Schmidt, Olga Chernomor, et al. 2020. “IQ-TREE 2: New Models and Efficient Methods for Phylogenetic Inference in the Genomic Era.” Molecular Biology and Evolution 37 (5): 1530–34. 10.1093/molbev/msaa015.

43. Mishima, Koichi, Tetsuro Watabe, Akira Saito, et al. 2007. “Prox1 Induces Lymphatic Endothelial Differentiation via Integrin Α9 and Other Signaling Cascades.” Molecular Biology of the Cell 18 (4): 1421–29. 10.1091/mbc.e06-09-0780.

44. Motta, Marialetizia, Giulia Fasano, Sina Gredy, et al. 2021. “SPRED2 Loss-of-Function Causes a Recessive Noonan Syndrome-like Phenotype.” The American Journal of Human Genetics 108 (11): 2112–29. 10.1016/j.ajhg.2021.09.007.

45. Murphy, Theresa L., Roxane Tussiwand, and Kenneth M. Murphy. 2013. “Specificity through Cooperation: BATF– IRF Interactions Control Immune-Regulatory Networks.” Nature Reviews Immunology 13 (7): 499–509. 10.1038/nri3470.

46. Nicenboim, J., G. Malkinson, T. Lupo, et al. 2015. “Lymphatic Vessels Arise from Specialized Angioblasts within a Venous Niche.” Nature 522 (7554): 56–61. 10.1038/nature14425.

47. Ocampo Daza, Daniel, and Tatjana Haitina. 2020. “Reconstruction of the Carbohydrate 6-O Sulfotransferase Gene Family Evolution in Vertebrates Reveals Novel Member, CHST16, Lost in Amniotes.” Genome Biology and Evolution 12 (7): 993–1012. 10.1093/gbe/evz274.

48. Oliver, Guillermo, Beatriz Sosa-Pineda, Sabine Geisendorf, Eric P. Spana, Chris Q. Doe, and Peter Gruss. 1993. “Prox 1, a *Prospero*-Related Homeobox Gene Expressed during Mouse Development.” Mechanisms of Development 44 (1): 3–16. 10.1016/0925-4773(93)90012-M.

49. Panara, Virginia, Hujun Yu, Di Peng, et al. 2024. “Multiple Cis-Regulatory Elements Control Prox1a Expression in Distinct Lymphatic Vascular Beds.” Development 151 (9): dev202525. 10.1242/dev.202525.

50. Papadogiannis, Vasileios, Dorit Hockman, Silvia Mercurio, et al. 2023. “Evolution of the Expression and Regulation of the Nuclear Hormone Receptor *ERR* Gene Family in the Chordate Lineage.” Developmental Biology 504 (December): 12–24. 10.1016/j.ydbio.2023.09.003.

51. Pedersen, Julia E., Christina A. Bergqvist, and Dan Larhammar. 2018. “Evolution of the Muscarinic Acetylcholine Receptors in Vertebrates.” New Research. eNeuro 5 (5). 10.1523/ENEURO.0340-18.2018.

52. Risebro, Catherine A., Richelle G. Searles, Athalie A. D. Melville, et al. 2009. “Prox1 Maintains Muscle Structure and Growth in the Developing Heart.” Development 136 (3): 495–505. 10.1242/dev.030007.

53. Rouse, Greg W., Nerida G. Wilson, Jose I. Carvajal, and Robert C. Vrijenhoek. 2016. “New Deep-Sea Species of Xenoturbella and the Position of Xenacoelomorpha.” Nature 530 (7588): 94–97. 10.1038/nature16545.

54. Sievers, Fabian, Andreas Wilm, David Dineen, et al. 2011. “Fast, Scalable Generation of High-quality Protein Multiple Sequence Alignments Using Clustal Omega.” Molecular Systems Biology 7 (1): 539. 10.1038/msb.2011.75.

55. Sosa-Pineda, Beatriz, Jeffrey T. Wigle, and Guillermo Oliver. 2000. “Hepatocyte Migration during Liver Development Requires Prox1.” Nature Genetics 25 (3): 254–55. 10.1038/76996.

56. Stollewerk, Angelika. 2016. “A Flexible Genetic Toolkit for Arthropod Neurogenesis.” Philosophical Transactions of the Royal Society B: Biological Sciences 371 (1685): 20150044. 10.1098/rstb.2015.0044.

57. Sutherland, Ben J. G., Thierry Gosselin, Eric Normandeau, et al. 2016. “Salmonid Chromosome Evolution as Revealed by a Novel Method for Comparing RADseq Linkage Maps.” Genome Biology and Evolution 8 (12): 3600–3617. 10.1093/gbe/evw262.

58. Tao, Shijie, Merlijn Witte, Robert J. Bryson-Richardson, Peter D. Currie, Benjamin M. Hogan, and Stefan Schulte- Merker. 2011. “Zebrafish Prox1b Mutants Develop a Lymphatic Vasculature, and Prox1b Does Not Specifically Mark Lymphatic Endothelial Cells.” PLOS ONE 6 (12): e28934. 10.1371/journal.pone.0028934.

59. Torii, Masa-aki, Fumio Matsuzaki, Noriko Osumi, et al. 1999. “Transcription Factors Mash-1 and Prox-1 Delineate Early Steps in Differentiation of Neural Stem Cells in the Developing Central Nervous System.” Development 126 (3): 443–56. 10.1242/dev.126.3.443.

60. Ungerer, Petra, Bo Joakim Eriksson, and Angelika Stollewerk. 2011. “Neurogenesis in the Water Flea *Daphnia Magna* (Crustacea, Branchiopoda) Suggests Different Mechanisms of Neuroblast Formation in Insects and Crustaceans.” *Developmental Biology*, Experimental and historical aspects of evolutionary bioscience, vol. 357 (1): 42–52. 10.1016/j.ydbio.2011.05.662.

61. Wang, Junfeng, Gamze Kilic, Muge Aydin, Zoe Burke, Guillermo Oliver, and Beatriz Sosa-Pineda. 2005. “Prox1 Activity Controls Pancreas Morphogenesis and Participates in the Production of ‘Secondary Transition’ Pancreatic Endocrine Cells.” Developmental Biology 286 (1): 182–94. 10.1016/j.ydbio.2005.07.021.

62. Weller, Mathias, and Diethard Tautz. 2003. “Prospero and Snail Expression during Spider Neurogenesis.” Development Genes and Evolution 213 (11): 554–66. 10.1007/s00427-003-0362-4.

63. Wigle, Jeffrey T., Kamal Chowdhury, Peter Gruss, and Guillermo Oliver. 1999. “Prox1 Function Is Crucial for Mouse Lens-Fibre Elongation.” Nature Genetics 21 (3): 318–22. 10.1038/6844.

64. Wigle, Jeffrey T., and Guillermo Oliver. 1999. “Prox1 Function Is Required for the Development of the Murine Lymphatic System.” Cell 98 (6): 769–78. 10.1016/S0092-8674(00)81511-1.

65. Xu, Peng, Xiaofeng Zhang, Xumin Wang, et al. 2014. “Genome Sequence and Genetic Diversity of the Common Carp, Cyprinus Carpio.” Nature Genetics 46 (11): 1212–19. 10.1038/ng.3098.

66. Zawisza-Álvarez, Michał, Claudia Pérez-Calles, Giacomo Gattoni, Jordi Garcia-Fernàndez, Èlia Benito-Gutiérrez, and Carlos Herrera-Úbeda. 2020. “The ADAR Family in Amphioxus: RNA Editing and Conserved Orthologous Site Predictions.” Genes 11 (12): 1440. 10.3390/genes11121440.

67. Zhang, Shujie, Ning Yu, Linfang Wang, et al. 2017. “Prox1 Represses IL-2 Gene Expression by Interacting with NFAT2.” Oncotarget 8 (41): 69422–34. 10.18632/oncotarget.17278.

68. Zhu, Denghui, Rong Huang, Peipei Fu, et al. 2019. “Investigating the Role of BATF3 in Grass Carp (Ctenopharyngodon Idella) Immune Modulation: A Fundamental Functional Analysis.” International Journal of Molecular Sciences 20 (7): 7. 10.3390/ijms20071687.

